# Statistical Methodology for Qualification of a Non-Clinical Risk Assessment Peptide:T Cell Proliferation Assay to Support Decision Making

**DOI:** 10.64898/2026.06.03.729894

**Authors:** Sophie Tourdot, Zhiping You, Andrew Ciarla, Rachel Hindin, Benjamin Keenan, Jennifer Calderini, Shea van den Broek, Christopher Lepsy, Timothy P. Hickling, Gregory Steeno

## Abstract

Antibody- and cell-mediated immune responses against biologics, should they occur, can impact treatment efficacy and potentially pose severe risks to patient safety. Therefore, developers have focused on advancing strategies to mitigate such unwanted immunogenicity. Opportunities to address immunogenicity early in the development process, particularly during the drug design phase, have been identified. *In vitro* and *in silico* tools that facilitate the identification and removal of sequence liabilities have been established. For example, human cell-based *in vitro* T cell assays can be used to identify and remove CD4+ T cell epitopes, which are known to play a critical role in the development of anti-drug antibodies against recombinant proteins products as well as the transgenes of gene and therapy. Despite their widespread use in the industry, most of these assays lack thorough characterization, which undermines confidence in the results and comparability across laboratories. In this study, concepts of immunogenicity bioanalytical assay validation for study design and analysis were applied to characterize an internal CD4+ T cell proliferation assay as fit-for-purpose. A statistical path was applied to establish data acceptance criteria for handling of replicates, positivity and negativity of a signal, and donor cohort size. A Bayesian analysis was also performed and is proposed as an approach for sequence de-risking decision making. The in-depth characterization of the CD4+ T cell proliferation assay described here allows for accurate interpretation of the assay outcomes, thereby enhancing confidence in using this approach for mitigating the immunogenicity of biologics by design.

## INTRODUCTION

Biologics have considerably improved the lives of patients across multiple diseases, including cancer, immunoinflammatory and metabolic disorders, as well as an increasing number of rare diseases. However, undesirable immune responses such as the development of anti-drug antibodies (ADA) and cytotoxic cellular responses to these drugs have the potential to compromise treatment efficacy and patient safety. (1-4) The probability to develop unwanted immunogenicity is associated with the aggregate risk of product, treatment, and patient-related factors (5, 6). Mitigation of immunogenicity and its consequences can be implemented in the clinic but ideally starts at the drug design stage through identification and reduction of the product’s intrinsic immunogenicity liabilities. This includes removal of sequence CD4+ T cell epitopes and post-translational modification sites, control of critical attributes including impurities content such as host cell protein and high molecular weight species, and of other biophysical properties that carry an immunogenicity risk, such as polyspecificity in the case of antibody formats or decoy receptors (7-9).

Application of a suite of *in silico* methods and *in vitro* assays has become a common strategy to identify and remove immunogenic CD4+ T cell epitopes from the primary structure of protein therapeutics and protein moieties of novel platforms. These include translated proteins of nucleic-acid, mRNA/LNP, or Adeno-Associated virus-based therapy products, as well as chimeric antigen receptors in cell therapy (10, 11). Non-clinical immunogenicity risk assessment (NCIRA) *in vitro* assays are of multiple formats and protocols, and unlike their bioanalytical assay counterparts, lack standardization (12, 13). In absence of harmonization, these assays might suffer from variable degrees of accuracy, a parameter difficult to quantify. Therefore, NCIRA *in vitro* assays undergo continuous development to increase confidence in their output (14-16).

Here, we report the characterization of a human peripheral blood mononuclear CD4+ T cell:peptide proliferation assay (PBMC:peptide assay) deployed at the drug design phase to identify potential immunogenic CD4+ T cell epitopes from the drug sequence (Figure 1). The objective of the study was to increase confidence in the risk assigned to a potential CD4 T cell epitope from the assay by establishing data-driven critical assay parameters including donor cohort size, donor response positivity to controls and test peptides thresholds. To this aim, a statistical path was applied to examine data integrity and mitigate the risk of mislabeling (false positive and false negative responses to peptides) at each step of the data processing workflow. Additionally, a new approach to peptide risk ranking was explored by means of a Bayesian analysis of peptide donor response rates, which provides a statistically based relative ranking that enables prioritization of CD4 T cell epitope de-risking.

**Figure 1.**
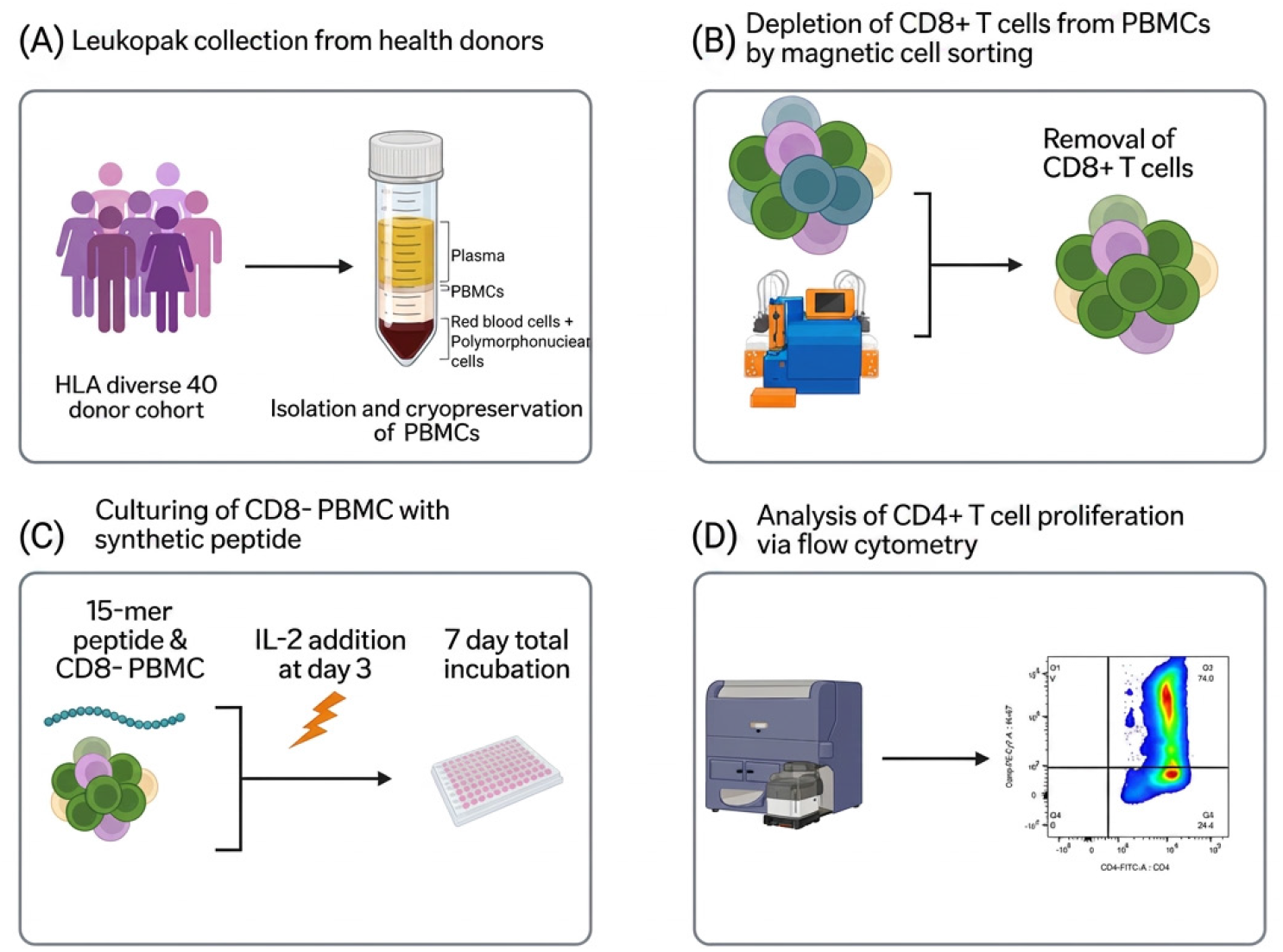
PBMC: peptide CD4 T cell proliferation assay principle. Peptide:T cell proliferation assay principle: Leukopaks from 40 HLA-diverse healthy donors are processed to isolate peripheral blood mononuclear cells (PBMCs) (A). PBMCs are thawed, rested, and CD8+ T cells removed (B). The remaining cells are incubated in six replicates with single 15-mer synthetic peptides with addition of IL-2 at day 3 (C); On day 7, proliferating CD4+ T cells are detected by flow cytometry (D).

This study suggests that applying a statistical path to cell-based *in vitro* NCIRA assays can increase data integrity and in return increase confidence in using these assays during drug design, mechanistic investigations or post-hoc analyses of clinical samples.

## MATERIAL AND METHODS

### Human peripheral blood mononuclear cells isolation and donor set selection

Leukopaks collected from HLA-typed healthy donors were purchased from StemCell Technologies (Vancouver, BC) and Charles River Laboratories (Wilmington, MA). Donor exclusion criteria included the use of nonsteroidal anti-inflammatory drugs (Advil, Motrin, Aleve & Aspirin), antihistamines (Zyrtec, Allegra, Claritin, Alavert, Azelastine & Benadryl) Pseudoephedrine, Mucinex, Afrin, all corticosteroids including nasal sprays (Flonase and Nasacort), and any immunomodulators (anti-inflammatory or oncology based), or recent known or suspected viral infection 8 weeks before blood draw. Peripheral blood mononuclear cells (PBMCs) were separated from leukopaks using either a Ficoll-Pacque gradient method (Cytiva Life Sciences, Marlborough, MA) or StraightFrom® Leukopak® human PBMC isolation kit (Miltenyi Biotec, Gaithersburg, MD) and the MultiMACS X separator. Briefly, for ficoll-pacque separation, leukopaks were diluted 1:1 with RPMI 1640 media (Cytivia Hyclone, Logan, UT), layered on ficoll-pacque, and centrifuged at 400g for 30min with no brake. The PBMC layer was collected, counted, washed twice with RPMI 1640 media, and frozen. Isolation from StraightFrom® Leukopak® human PBMC kits was performed by the manufacturing instructions using the MultiMACS X separator. Once the PBMC were isolated they were counted, washed twice with autoMACS® Rinsing Solution supplemented with 0.05% BSA, (Miltenyi Biotec, Gaithersburg, MD) and frozen. PBMCs were cryopreserved in fetal bovine serum (Cytiva Life Sciences, Marlborough, MA) and 10% dimethyl sulfoxide (DMSO, Sigma-Aldrich, St. Louis, Missouri) immediately after isolation and within 24 hours of Leukopak collection at a controlled rate of - 1°C per minute in a -80°C freezer using CoolCell freezing containers (Corning Life Sciences, Tewksbury, MA). PBMCs were then transferred to long-term storage at -190°C in vapor-phase liquid nitrogen. If donors HLA genotypes were not provided by leukopak vendors (Charles River Laboratories and StemCell Technologies), RNA was extracted from frozen PBMCs using DNeasy Blood & Tissue Kits (Qiagen, Germantown, MD) and analyzed by LabCorp (Burlington, NC). Study donor sets were selected using an internal algorithm designed to match HLA-DRB1 allele frequency distribution to that of the North American population.

### Test peptides and controls

All 15-mer peptides were synthetized by WuXi AppTec, Wuxi, Jiangsu, China, with an acceptance criterion of ≥95% purity. Class II–associated Ii chain peptide (CLIP, (17)) and HLA class II Cytomegalovirus, Epstein-Barr virus, Influenza virus, and Tetanus toxin peptide pool (CEFT, Cellular Technology, Shaker Heights, OH) were used as low and high donor response frequency sensitivity controls respectively. Staphylococcal enterotoxin Type B (SEB) (Toxin Technologies, Sarasota, FL) was used as an assay system control at a final concentration of 250 ng/mL. All test peptides and controls were added to cells at a final concentration of 5 µM.

### PBMC:peptide T cell proliferation assay

CD4+ T cell responses to synthetic peptides were measured using an *in vitro* T cell proliferation assay. Cryopreserved PBMC from healthy donors were thawed, washed in AIMV media (Thermo Fisher Scientific, Waltham, MA), counted (Cellometer SD100 cell counter, Nexcelom Bioscience, Lawrence, MA), and incubated on ice for 50 minutes. Following incubation, cells were centrifuged and washed in 40mL MACS Separation Buffer (autoMACS® Rinsing Solution supplemented with 0.05% BSA, Miltenyi Biotec, Gaithersburg, MD). Human anti-CD8 Microbeads (Miltenyi Biotec, Gaithersburg, MD) were added to the PBMCs according to the manufacturer instructions. Labeled cells were washed once and resuspended in MACS Separation Buffer and loaded onto an AutoMACS® Pro Separator (Miltentyi Biotec, Gaithersburg, MD) to allow for the depletion of CD8+ cells according to the manufacturer’s protocol. The CD8+ T cell depleted PBMC was washed twice in AIMV media and seeded at 5.0 × 10^5^ cells per well into 96-well round-bottom plates (Corning Life Sciences, Tewksbury, MA) in replicates of 6. Lyophilized synthetic test and control peptides were reconstituted in DMSO and added to the wells at a final concentration of 5µM per well. Peptide-treated cells were cultured at 37°C and 5% CO_2_ for 3 days. On Day 3, cultures were supplemented with 5 ng/mL of recombinant human IL-2 (Sigma-Aldrich, Burlington, MA). Cells were cultured at 37°C and 5% CO_2_ for an additional 4 days before immunostaining and acquisition via flow cytometry.

### Flow cytometry

After 7 days of cell culture, plates were centrifuged at 500g for 5min, and the supernatant discarded. The cell pellets were washed with BD Pharmingen™ Stain Buffer with 2% FBS (BD Biosciences, Franklin Lakes, NJ) and then surface stained for CD3 (PE, clone UCHT1, Biolegend, San Diego, CA), CD4 (FITC, clone SK3, Biolegend, San Diego, CA), and live/dead (near-IR, Thermo Fisher Scientific, Waltham, MA) for 30min at 4°C. Following incubation, cells were washed with stain buffer, fixed and permeabilized using a Foxp3/Transcription Factor Staining Buffer Set (Thermo Fisher Scientific, Waltham, MA) for 30min at 4°C. The cells were then washed with permeabilization buffer and intracellularly stained for the proliferation marker Ki-67 (PE-Cy7, clone 20Raj1, Thermo Fisher Scientific, Waltham, MA) for 30min at 4°C. The cells were then washed twice with stain buffer and 10,000 CD3+/CD4+ T cell events were acquired within 48h using BD FACSCelesta™ or BD LSRFortessa™ flow cytometers (BD Biosciences, Franklin Lakes, NJ). Raw data were analyzed using BD FlowJo software.

### Statistical Analyses

Experimental design combined with both parametric and nonparametric statistical summarization, modeling methods, and simulation were applied to assess assay performance, replication strategy, critical quality and response thresholds, donor cohort size, and peptide differentiability.

Data spanning formal qualification and early portfolio support were consolidated to quantify different variability sources representing repeatability and reproducibility. Collectively, data were generated from 5 screens, 131 peptides, 23 different runs, 163 donors, each with a target of six replicates per donor:peptide combination. Non-representative peptides and cell functionality controls were removed before the variance analysis. Final data volume for statistical modeling was 25731 observations. Classical linear mixed effect models (18) were employed to estimate variance sources and goodness-of-fit diagnostics were performed to assess distributional assumptions. The linear mixed effects model is expressed as

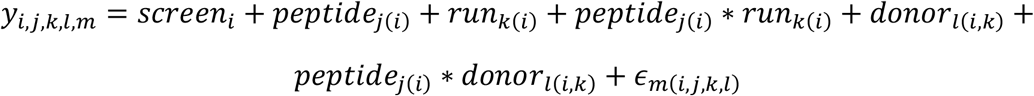

where *y*_*i,j,k,l,m*_ is the CD4^+^ T-cell proliferative response within the *i*^*th*^ screen, the *j*^*th*^ peptide, the *k*^*th*^ run, the *l*^*th*^ donor, and the *m*^*th*^ replicate of the screen-peptide-run-donor combination. The hierarchical model includes nested random effects, such as donors within a screen:run and crossed random effects, *peptide*run* and *peptide*donor*, interpreted as heterogeneous peptide random effects across runs (presumed negligible) or across donors (presumed non-negligible). The residual term, *ϵ*, represents well-to-well replicate variability to infer precision of summary measures as a function of replicate number. Distributional assumptions were graphically assessed via QQ-plots.

To reduce sensitivity to anomalous observations and heavier-tailed distributions, robust measures of location and spread were applied to identify acceptable replicate and aid in defining donor positivity. As a quality control check, replicate dispersion for each donor:peptide combination is evaluated using a robust coefficient of variation, defined as the sample median absolute deviation (MAD) divided by the sample median. If acceptable, the median 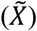 is used as the measure of T-cell proliferation. Definitions of the sample median and sample *MAD* are expressed below, where 1) *X*_(_*i*_)_ is the *i*^*th*^ order statistic in the median definition, and 2) 1.4826 is a scaler to make the estimate consistent under normality.

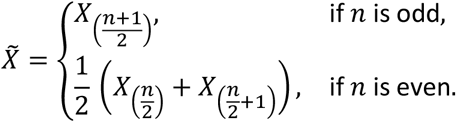

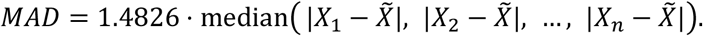

For any donor:peptide combination, the stimulation above background is defined as the difference between the median T-cell proliferative response with the peptide minus the median baseline response control; 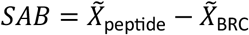. The donor is considered a positive response for that peptide if the SAB is both greater in magnitude than a threshold, *c*^*+*^, and demonstrates statistical significance via a robust 1-*α* level *t*-test.

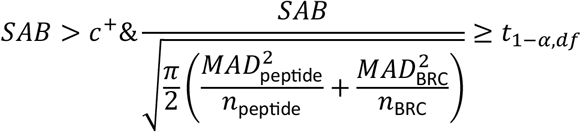

For each peptide-donor combination, robust residuals, 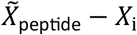, were calculated, stored, and bootstrapped with replacement to establish SAB quantities of interest: (1) the donor positivity threshold, *c*^+^, for a targeted specificity, and (2) a lower quality limit, *c*^−^, defining excessive baseline response control. For both objectives we assume the true SAB is zero, implying no additional T-cell proliferation due to the presence of the peptide. Under that assumption, *n = 6* residuals for both a virtual peptide and virtual baseline response are drawn with replacement 100000 times, and the SAB from the two medians is calculated and stored. The simulated SAB distribution is visualized and SAB quantities of interest under the scenario where the peptide has no risk are inferred.

Peptide donor response rates (*θ*) were analyzed using a beta–binomial Bayesian model. The number of positive responding donors for a given peptide was modeled using a binomial likelihood, *p*(*y*|*θ*), with the underlying probability of donor response rate assigned a beta prior distribution, *p*(*θ*). Posterior inference was obtained by combining the likelihood of the observed data with the prior, scaled by the marginal *p*(*y*).

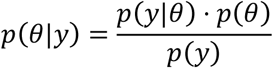

The benefits in using this statistical path include 1) incorporation of prior donor-response rate knowledge, 2) efficient use of smaller, possibly unbalanced, data sets and 3) direct probability statements on peptide ranking. For (1), prior knowledge could vary from weak / non-informative, e.g., a new screening peptide, or strong / informative, e.g., for a control peptide. The donor response rate for a peptide is inferred by the mean or median of the posterior distribution with associated (1-*α*)*100% credible intervals. To compare response rate for peptides *P*_*i*_ and *P*_*j*_, the probability that *P*_*i*_ has a lower donor response rate than *P*_*j*_ is estimated as Pr(*θ*_*i*_ *< θ*_*j*_ | prior, data) using Monte Carlo sampling from the respective posterior distributions.

Simulation was performed to comprehend the impact of donor cohort size on successful rank-ordering of peptides. For peptides *i* and *j* with true donor response probabilities *θ*_*i*_ and *θ*_*j*_, *n*_*d*_ donor-level latent responses were simulated as *y*_*kℓ*_ ∼ Bernoulli(*θ*_*k*_) (*k* ∈ {*i,j*}, *ℓ*=1,…, *n*_*d*_). A new variable was created, *z*_*kℓ*_, such that positives (*y*_*kℓ*_ = 1) were mapped to a continuous assay response *μ* with *μ* > *c*^*+*^, and negatives (*y*_*kℓ*_ = 0) to 0. Independent measurement error was added: 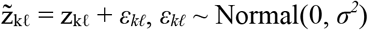. Observed binary responses were regenerated via thresholding 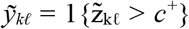, yielding counts 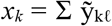. With uniform priors *θ*_*k*_ ∼ Beta(1,1), posteriors were *θ*_*k*_ | data ∼ Beta(1 + *x*_*k*_, 1 + *n*_*d*_ − *x*_*k*_). The rank-order probability Pr(*θ*_*i*_ < *θ*_*j*_ | data) was estimated by Monte Carlo as the proportion of M=10,000 paired posterior draws where *θ*_*j*_ > *θ*_*i*_.

All analyses and visualizations were completed in both R 4.5.1 and SAS 9.4.

## RESULTS

The PBMC:peptide assay was developed internally to guide immunogenicity sequence de-risking during drug design. The assay is applied to assess the capacity of CD4+ T cell epitopes predicted by *in silico* methods to stimulate a T cell response and thus constitute potential sequence liabilities. The T cell risk of a given epitope is expressed as a percentage of donors that respond to that epitope in a study, or donor response rate. Donor response rates can be compared to prioritize protein engineering to remove epitopes within the pool of risky epitopes. Before starting the assay qualification, terms and definitions were established to describe the statistical path with clarity. It was particularly important to differentiate responses at the donor level, i.e. the response of a given donor to a given peptide or “donor:peptide pair response” from the response at the cohort level, i.e. the total number of donors that respond to a given peptide or “peptide donor response rate” **(Table 1)**.

**Table 1.** Terms and definitions.

| Term | Definition |
| --- | --- |
| **Donor set/cohort** | Group of donors included in a run |
| **Run** | One independent PBMC assay experiment conducted with one set of donors |
| **Donor:peptide pair** | Response of an individual donor to a given peptide |
| **Donor set response rate** | Number of positive donor:peptide pairs in a run for a given peptide |
| **Stimulation above background (SAB)** | Difference between the signal observed in response to a test article (peptides, controls) and the signal observed in absence of stimulus |

### 1. QUALIFICATION

#### 1.1. Inclusion of assay controls

The accuracy of a procedure can be improved by establishing reference points. During assay development the reference points, or controls, indicate whether the reaction conditions are optimal, help identify potential issues and guide the establishment of data quality checks. Once the assay is established, controls serve as acceptance/exclusion criteria at each step of the data analysis process, leading to the final decision-making output. At the run level, a discrepancy between observed and expected donor response rates to the High or Low Sensitivity Control (HSC, LSC, respectively) signals a technical issue; the run will be rejected. At the donor level, donors that i) do not respond to a non-antigen specific, pan-T cell activator compound (Cell Functionality Control, CFC), ii) exhibit a higher response in absence of stimulus (Baseline Response Control, BRC) than in presence of peptides, or iii) respond to all test peptides included in a run will (i) or might (ii, iii) be excluded from the final donor response rate analysis. The nature of the controls used in the assay are detailed in **Table 2**. The rational for donor exclusion/inclusion in the three categories is provided at the “Data integrity” and “Peptide donor response rates sections”.

**Table 2.**
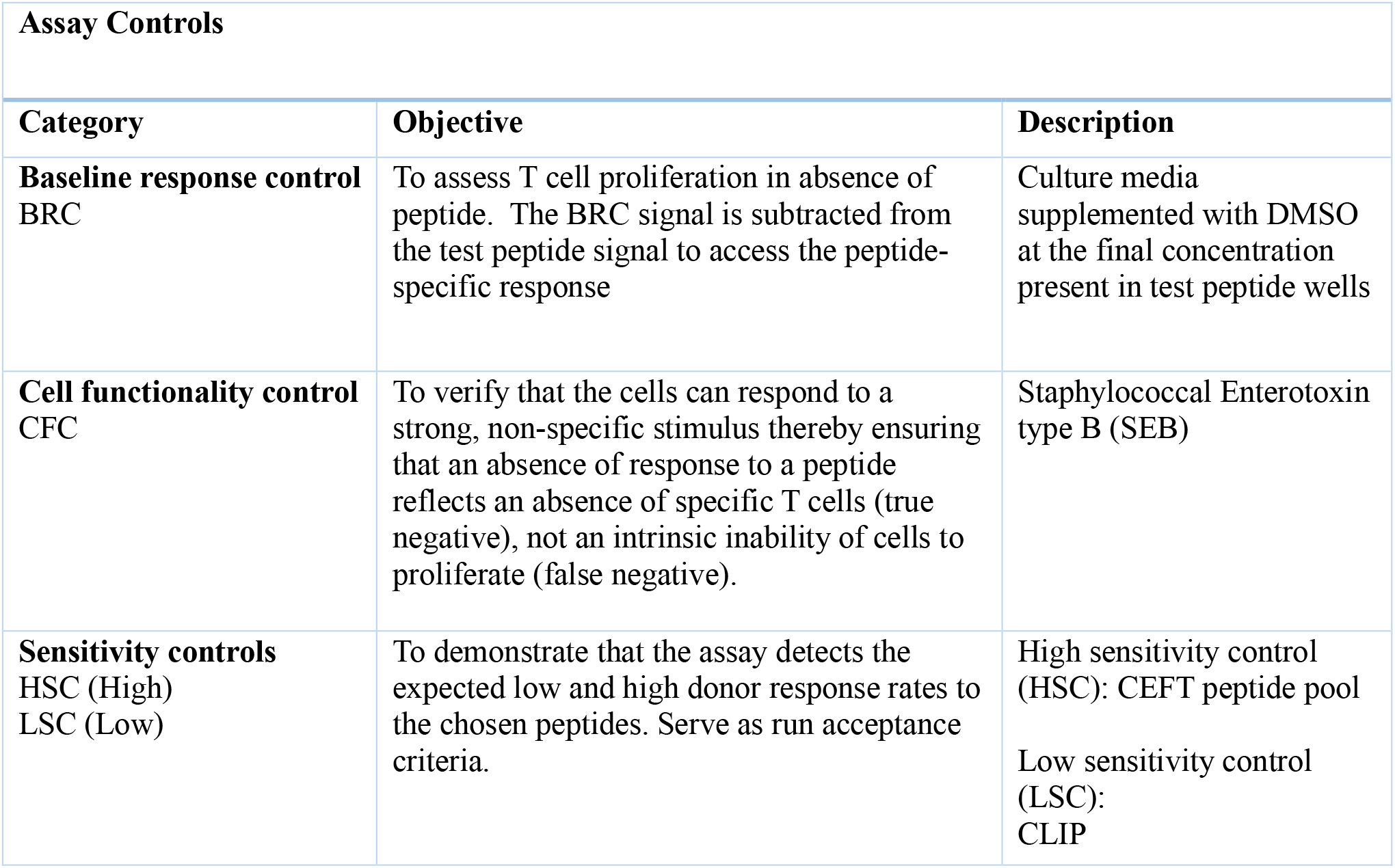
Controls used in the assay. The assay controls categories, objectives and terminology follow the European Immunogenicity Platform best practices recommendations (16).

#### 1.2. Variability of the assay

A fit for purpose qualification of the assay was performed to assess the precision of the method. To this end, a variance components analysis was conducted to quantify the respective contributions of known sources, including operators, instruments, well replicates, reagents, donors, and day of assay execution, to the overall assay variability. For simplification and interpretability, individual assay influences were grouped into method sources of variability, such as run-to-run and within-run effects, and non-method sources, such as screen, peptide, and donor effects. Where warranted, specific factors (e.g., operator effects) were interrogated by modeling them as random effects nested within the run structure and evaluating their associated variance components relative to total assay variability, thereby assessing the extent to which operator to operator differences contributed meaningfully to overall assay precision. The design structure of the designated qualifications study is shown in **Figure 2(A)**. Data from sequence de-risking of internal programs was added to the data set to inform the previously expressed linear model. Variance contributions for both the method and non-method sources are illustrated in **Figure 2(B)**. For the method sources, run-to-run is the highest source of variability, which is a composite of different operators, days, etc. The run*peptide term is small and is interpreted as consistent deviations across the runs regardless of peptide. The residual term is the replicate variability and *n = 6* replicates results in a standard-error of the mean around 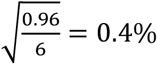 in magnitude, or about 10% CV for typical baseline control responses. For the non-method sources, by design the donor-to-donor component is the highest source. The peptide*donor is a bit larger and simply indicates T-cell proliferation of a peptide is a function of donor. Finally in **Figure 2(C)**, residual diagnostics from the normal QQ-plot inform that the replicate distribution is heavy tailed and that reliable inferences on peptide risk would greatly benefit from robust statistical measures as done here.

**Figure 2:**
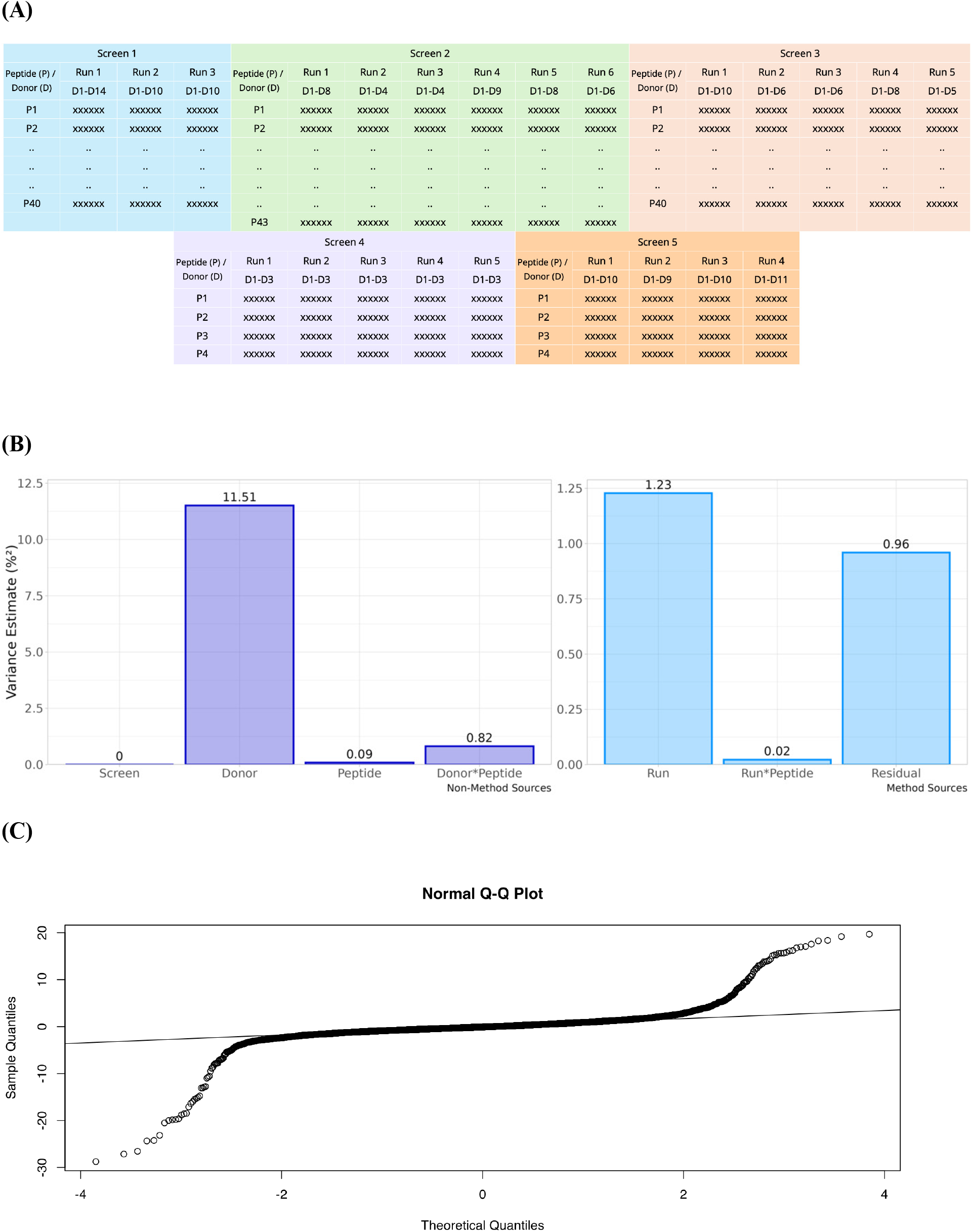
Variance Analysis. Variance Component Design and Analysis. (A) Design structure from collected data that informs linear mixed effects model. (B) Variance estimates across non-method and method sources. (C) Normal QQ-plot of residuals indicating much heavier tailed distribution and the need for robust estimation.

#### 1.3. Critical parameters

##### 1.3.1. Response signal

A donor response to a peptide or control is expressed as a stimulation above background (SAB). In this assay, SAB is calculated by subtracting the BRC percentage of T cell proliferation from that of the peptide or control. Donor:peptide or donor:control pair responses will be considered positive if they reached an established threshold. To establish the SAB positivity threshold, a simulation of SAB variation around zero for an expected donor:peptide pair negative response was conducted using the empirical residuals as described above. **Figure 3** depicts the SAB positivity cut-off, *c*^*+*^, calculated to provide a targeted (1-*α*)*100% specificity that a truly negative peptide would produce SAB values below *c*^*+*^. For our application, we targeted 95%, or a 5% false positive rate, resulting in *c*^*+*^ *= 0*.*8*. Additionally, because SAB is expressed as a signal difference, peptide:donor pairs can exhibit negative SAB values when intrinsic method variability causes a donor’s response to test peptides lower than the response to the BRC. However, very negative observed SAB values are a function of a disproportionately high background response, which might signal a donor-related issue. Hence, considering those donors as non-responders and including it in the final analysis carries the risk of an inaccurate peptide donor rate. To remediate the risk of donor:peptide pair response mislabeling, the same simulation was used to determine a lower limit of SAB negativity. The lower quality limit was calculated at 99.9%, where values below this limit occur 0.1% of the time under normal assay conditions. Based on our simulation, *c*^*-*^ was set at -2.5.

**Figure 3:**
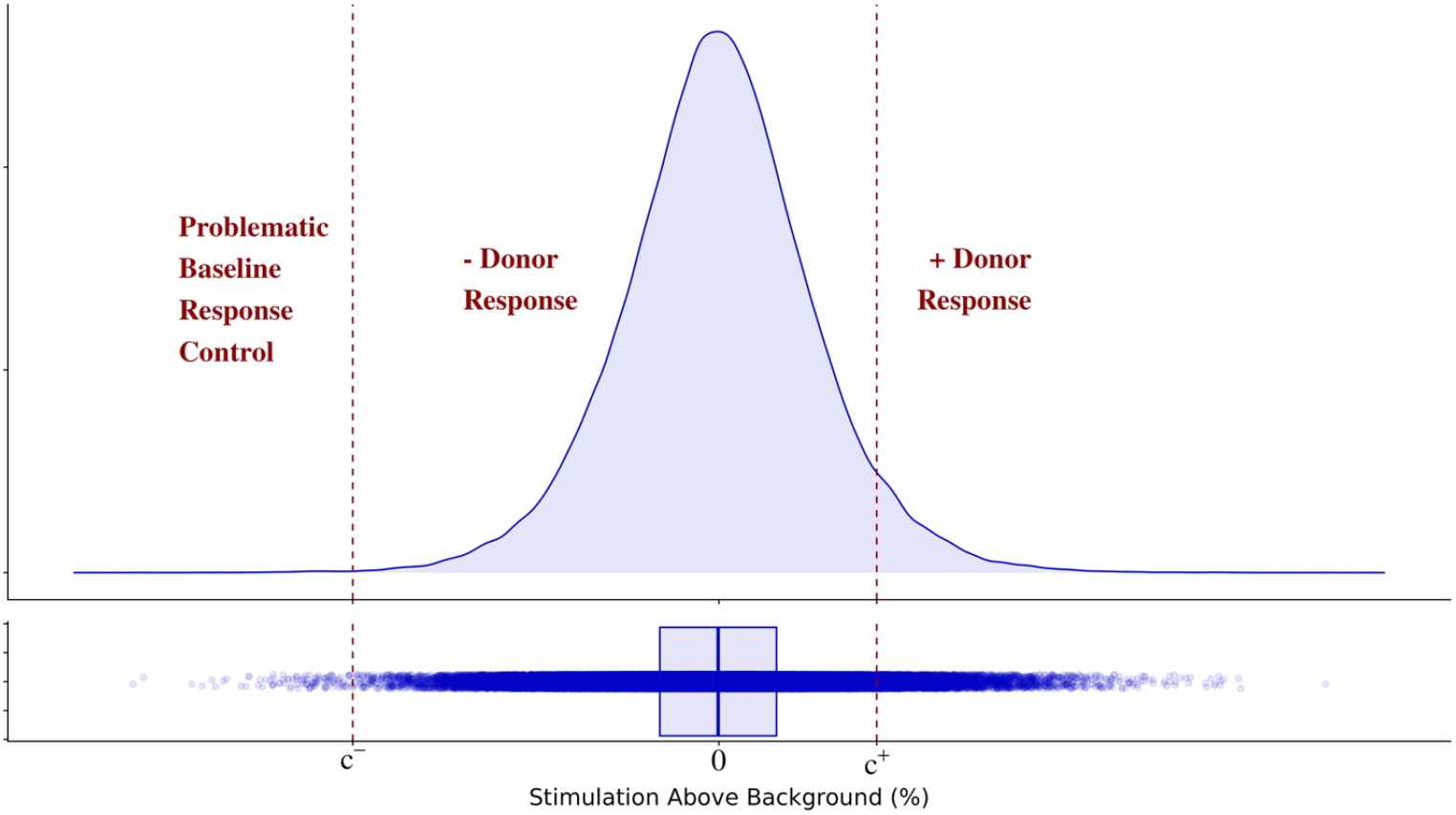
Stimulation above background positivity and true negativity cut-offs. Bootstrapped distribution of *SAB* values from *n* = 6 replicates under the assumption that the tested peptide has zero effect on T-cell proliferation. Calculated critical values are a lower 99.9% control limit that defines excessive baseline response control (*SAB* < *c*^*-*^) and an upper 95% specificity limit that defines a positive donor response (*SAB* > *c*^*+*^).

##### 1.3.2. CD4+ T cell count

When running the assay, the number of cells distributed in each well is based on total CD8+ T cell-depleted PBMCs. Therefore, donor-to-donor variability in the number of CD4+ T cells surveyed is expected. However, too low a number of CD4+ T cells might lead to no or a low frequency of CD4+ T cell precursors able to recognize peptides and prevent the detection of peptide-specific responses, leading to mislabeling donor:peptide or donor:control responses as negative responses. To investigate the impact of CD4+ T cell content variability, a PBMC assay including samples with or without correction for CD4+ T cells content in the same run was performed (data not shown). Data analysis showed that the SAB variability between the two methods near the positive cut-off is very comparable, leading to equivalent assay specificity and sensitivity. We did observe higher SAB variability with the total CD8+ T cell-depleted PBMCs when corresponding SABs were (much) higher than the positive threshold. However, this would not impact classifying a donor as a positive responder.

##### 1.1.1. Donor cohort size

Next, the number of donors included in a run to correctly rank-order peptide donor response rates with high success was investigated. The minimum true donor response rate difference between peptides *P*_*i*_ and *P*_*j*_ was targeted at *θ*_*i*_ *– θ*_*j*_ = 15%. For example, (*P*_*i*_, *P*_*j*_) combinations include (20%, 5%), (25%, 10%), …, (50%, 35%). The donor cohort size was interrogated by performing a simulation of the Bayesian analysis path across the various true donor response rate differences with donor cohort sizes ranging from 10 – 60, then calculating the median rank-order probability, akin to a power analysis. As shown in **Figure 4**, 40 donors are needed to correctly rank order peptides with a true 15% response rate difference at least 80% of the time (median rank-order probability > 0.8). From the same figure, a true 20% response rate difference in peptides is correctly rank ordered > 85% of the time. The assay has capacity for 40 donors, thus no additional impact on throughput.

**Figure 4:**
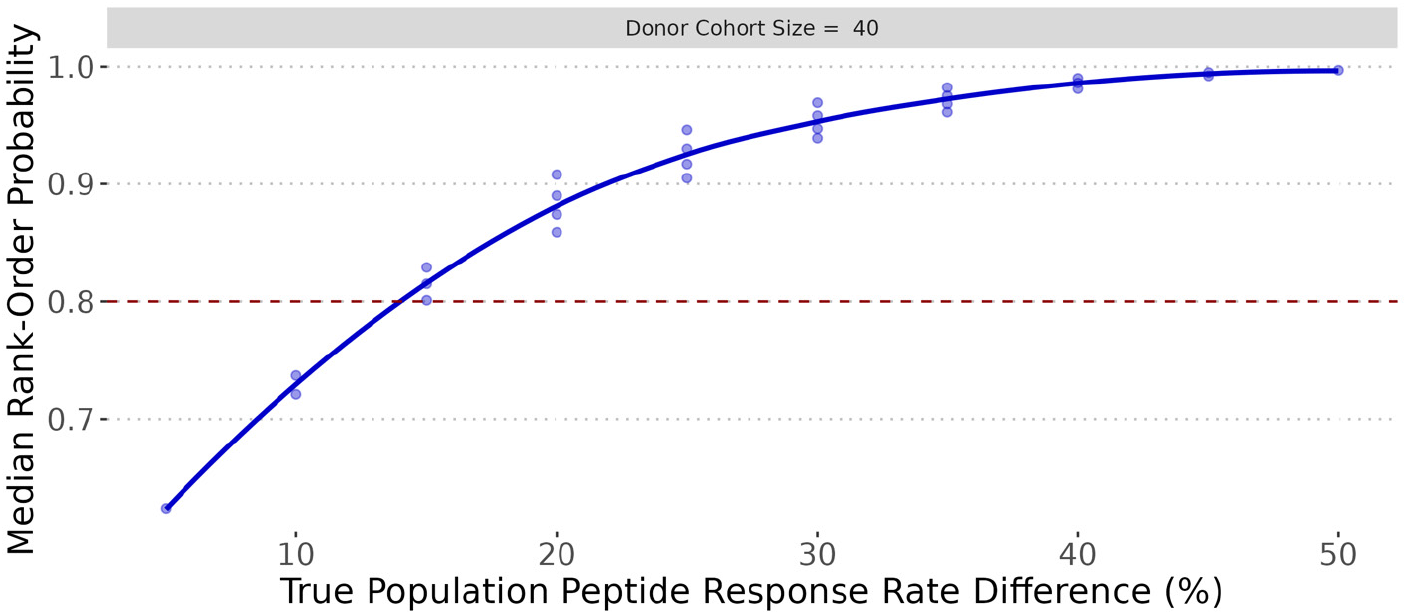
Donor cohort size. Simulated rank-order probabilities between two peptides as a function of the true difference in donor response rate. Different symbols represent various true donor response rates for each peptide that map to a specific difference. E.g., 15% donor response rate difference occurs if peptide one (P1) has true 25% donor response rate and the peptide two (P2) has 10%. Likewise, P1 has 30%, P2 has 15%, etc). Superimposed is LOESS smoother indicating that median rank order probability > 0.8 (80%) if true donor response rate difference P1-P2 = 15%.

### 2. DATA INTEGRITY

Having established the performance of the assay and determined the SAB positivity, negativity cut-offs and threshold, as well as the donor cohort size, a thorough interrogation of the data analysis workflow leading to the decision-making output was conducted. The aim of the investigation was to identify and remediate the risk of data misinterpretation at each step of the process.

#### 2.1. Replicate dispersion

The first data processing step is the conversion of the 6 replicates individual values into a single value representative of the donor:peptide pair response. As shown in **Figure 5**, the replicates exhibit variable levels of dispersion with discrete patterns. The use of the mean to access the replicate central value is common practice in cell-based assay. Because of the heavy-tailed distribution shown in Figure 2(C), using the mean has the potential to overestimate (e.g., Peptide 2 from Donor 1) or underestimate of the replicate central value and in return, generate false negative and false positive SABs. In contrast, because it is resistant to extreme values, the median is a robust measure providing a more accurate replicate central value representing a typical response. However, when the 6 replicates form 2 distinct and distant groups of 3 (e.g., Peptide 4 from Donor 1), the central value provided by the median (or mean) inaccurately reflects the response observed. To remediate this risk, a data integrity check was implemented by calculating the robust replicate CV, a variability measure resistant to outliers yet sensitive enough to reject anomalous data. If data from a donor:peptide set are above a predetermined threshold (e.g., 30%), those will be excluded from the donor response rate calculation. As shown in Figure 5, adding the robust CV as an exclusion criterion correctly rejected data from Peptide 4, but data from Peptide 2 data were determined acceptable in the presence of one discordant observation.

**Figure 5:**
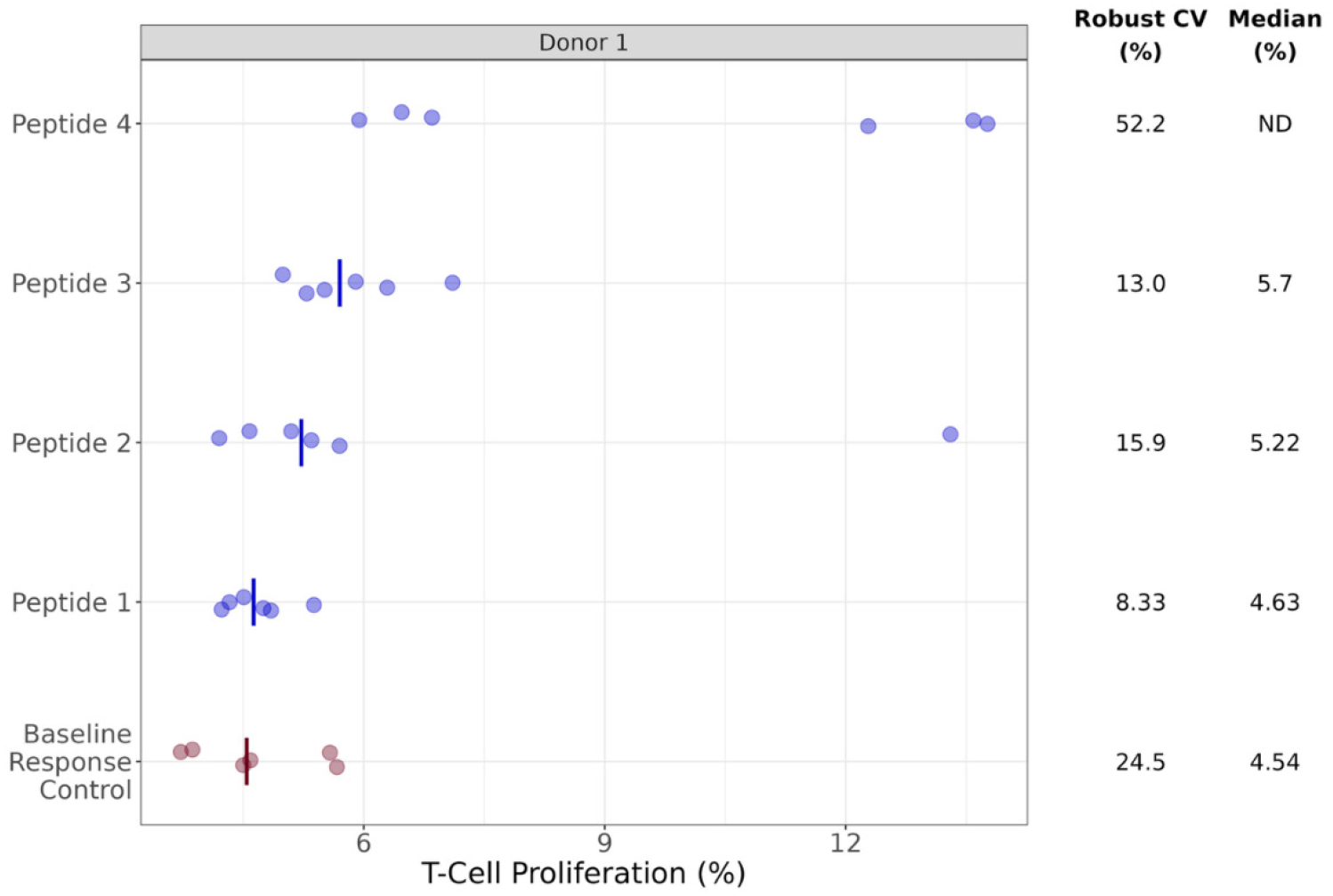
Replicate Dispersion and Central Value. Distribution of Donor 1 data (·) across peptides and baseline response control with superimposed median (|), if applicable based on acceptable robust coefficient of variation (< 30%). Peptide 2 data pass, even in the presence of one discordant observation. Peptide 4 data fail with the two separate observation groups. ND = Not Determined.

#### 2.2. Donor:peptide pair response positivity

Commonly, whether SAB is expressed as a fraction or a difference, a donor:peptide pair response is considered positive on the sole criteria of SAB being equal or superior to an established positivity threshold. In this approach, no attention is paid to the magnitude of the difference between the BRC and peptide/control T cell proliferation relative to the variability. However, very low BRC values can create artificially positive SABs: the smallest increase of T cell proliferation in the test conditions will pass the SAB positivity threshold although that percentage is not biologically significant. Similarly, very high SAB values might occur due to small samples and high variability, even with robust measures. To protect against the risk of mislabeling (false negative and positive donor:peptide pair responses), the response positivity definition was revised to incorporate the magnitude of the difference between the BRC and test peptide signals relative to the observed variability. A donor:peptide pair response will be labeled positive only if the SAB is equal or superior to the SAB positivity threshold determined during the qualification and the difference in T cell proliferation between the BRC and the test condition is statistically significant. Using the equation above, donor positivity is inferred if *SAB > c*^*+*^ *= 0*.*8* and the robust t-statistic > a critical t-value at a pre-specified significance level of *α*.

#### 2.3. Inclusion of donor:peptide pairs in the calculation of peptide donor response rate

After implementing data integrity measures for raw data processing and donor:peptide pair response labeling, the criteria required for including a donor:peptide pair in the calculation of the peptide response rate, which informs decision-making, were examined.

##### 2.3.1. Magnitude of the response to the cell functionality control

Healthy T cells are expected to respond to a non-antigen-specific strong stimulus. By assessing this ability for each donor, the CFC ensures that when a donor exhibits negative responses to a peptide or sensitivity control, those are due to an absence of specific responding T cells (true negatives) and not to an intrinsic inability of cells to respond (false negatives). Therefore, donors that do not satisfactorily respond to the CFC will be excluded from the donor response rates calculation. As an example, **Figure 6(A)** shows Donor 3 not appearing to sufficiently respond to the CFC control. To be applicable to any study, the CFC SAB positivity cut-off should integrate a component of assay variability. For this, the distribution of CFC SABs across all donors in the data set was examined. As illustrated in **Figure 6(B)** > 80% of the time, a donor’s CFC SAB will exceed 20%. To keep a reasonable donor rejection rate, the CFC SAB positivity cut-off is established at the lower 10 percentile, which equates to an SAB = 12%. And donors that have a CFC SAB less than that are rejected. Because 1 in 10 donors per run are expected to fail the CFC, additional donors are added to each study to remain within a cohort size range that powers the assay to satisfactorily differentiate donor response rates.

**Figure 6.**
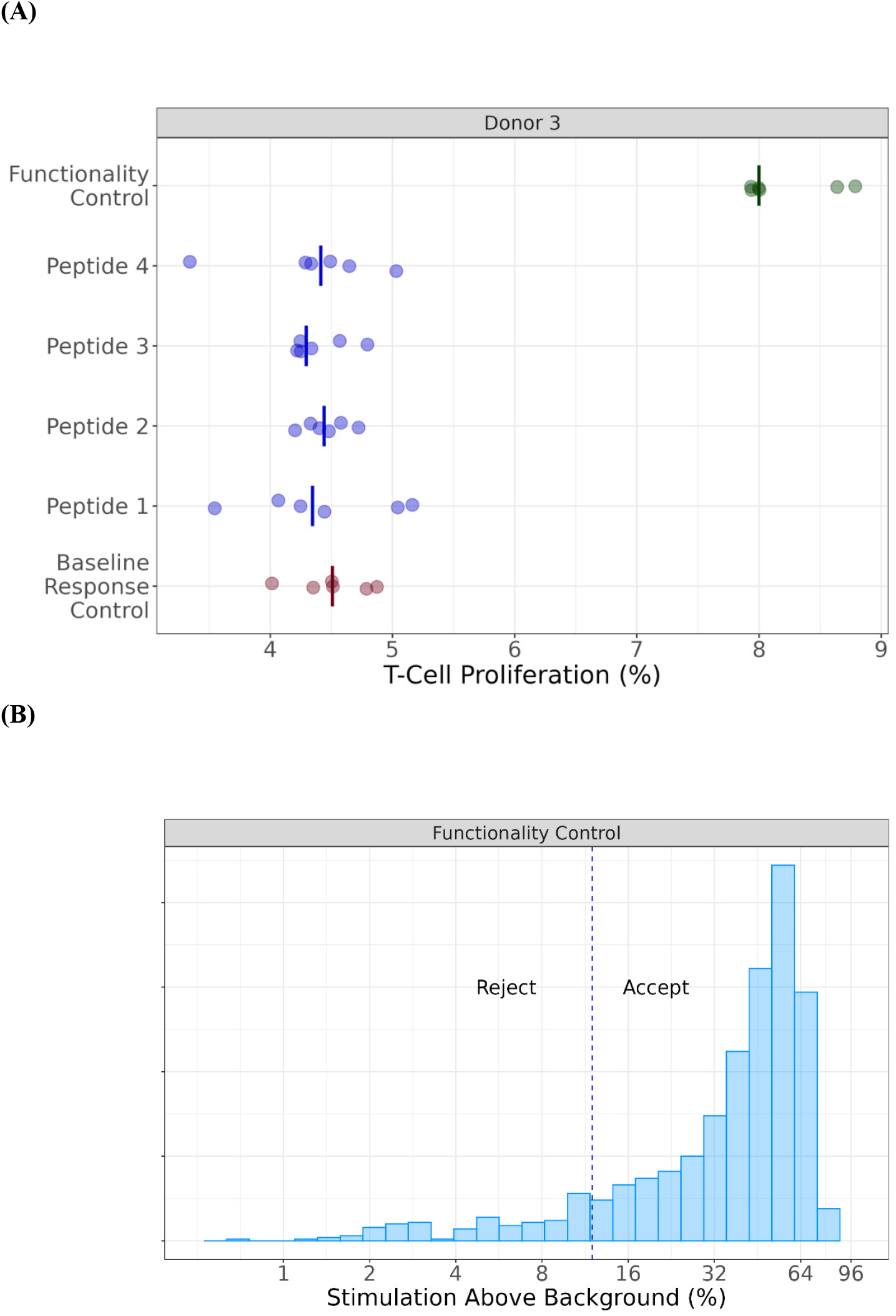
Cell Functionality Control Performance. (A) Distribution of Donor 3 data (·) across cell functionality control, peptides, and baseline response control with superimposed median (|). CFC data and median for this donor are low; observed Stimulation Above Background (SAB) is 8 – 4.5 = 3.5. (B) Empirical distribution of Cell Functionality Control Stimulation Above Background (SAB) values with superimposed reference line at the lower 90% quantile where SAB = 12%. Within an experiment, donors that fall below that threshold are not included in data analysis.

##### 2.3.2. Response to all test peptides

A prerequisite for peptides to be presented by HLA molecules for recognition by T cells is for the sequence to contain a defined HLA allele-specific binding motif. Thus, in theory, the likelihood of a donor presenting and responding to the entire set of peptides included in a study is very low. It implies that the observed positive responses might not be genuine, but rather due to technical issues with the donor. To mitigate the risk of false donor:peptide positive responses to some or all peptides tested, the donor might be excluded from the final analysis. However, considering that multiple HLA alleles share the same binding motif (HLA supertypes and that peptides can display some degree of promiscuity (19), the probability of these responses to be real is not null. Indeed, while positive responses to an entire set of 40 peptides highly diverse in sequence and HLA binding motifs have a low probability of being genuine, positive responses to a much smaller set, or a set of peptides that have a high degree of homology might be biologically pertinent. As no statistical method was deemed appropriate for establishing the upper limit of positive responses considered genuine, each case was evaluated on an individual basis. Decisions were made based on both the number and characteristics of the peptides evaluated in the study.

### 3. TEST ARTICLE RISK RANKING

After accounting for potential mislabeling of donor:peptide response pairs, we totaled positive responses for each peptide and calculated the donor response rate—the count of positive donor:peptide pairs releative all donors tested per peptide. The decision to eliminate a CD4+ T cell epitope during molecular design is determined by the observed donor response rate; peptides that elicit responses from a greater number of donors are regarded as presenting higher risk. Most commonly, the risk is assigned by grouping donor response rates in discrete risk categories. For instance, peptides with a donor response rate up to 10%, 20% or 50% might be classified as low, medium and high risk, respectively. As the lower and upper limits of each category are frequently based on a laboratory’s historical methods or common practices in the field, an alternative, data-driven approach to express an epitope T cell risk was investigated. The Bayesian analysis path consists of incorporating prior knowledge, which might be weak or strong, plus newly observed data to update our belief (20) and provide project teams a figure of merit to differentiate the investigated peptides. As peptides included in a study would not have been tested before in the assay, the prior knowledge applied to test peptide data was the expectation that the donor response rate is comprised between 0 and 1, i.e., the response rate cannot be negative nor greater than 1. By contrast, because HSC and LSC are included in all studies, a large body of donor response rates to both controls had been generated and constituted prior knowledge; representative examples of both cases are shown in **Figure 7(A)**. The output of the analysis is the expression of a peptide risk as a probability to be less, equally or riskier than other peptides, including LSC and HSC, given the data in the study and prior knowledge. As illustrated in **Figure 7(B)**, peptide 2 (P2, ∼12% predicted donor response rate) has an 86% chance to be less risky than peptide 1 (P1, ∼24% predicted donor response rate). As there are multiple tested peptides in a typical screen, this is easily extended and visualized to all possible pairwise comparisons as shown in **Figure 7(C)**. For example, peptide 3 (P8, 19.4% predicted donor response rate) has only 71% chance to be riskier than peptide 5 (P9, 13.8% predicted donor response rate).

**Figure 7:**
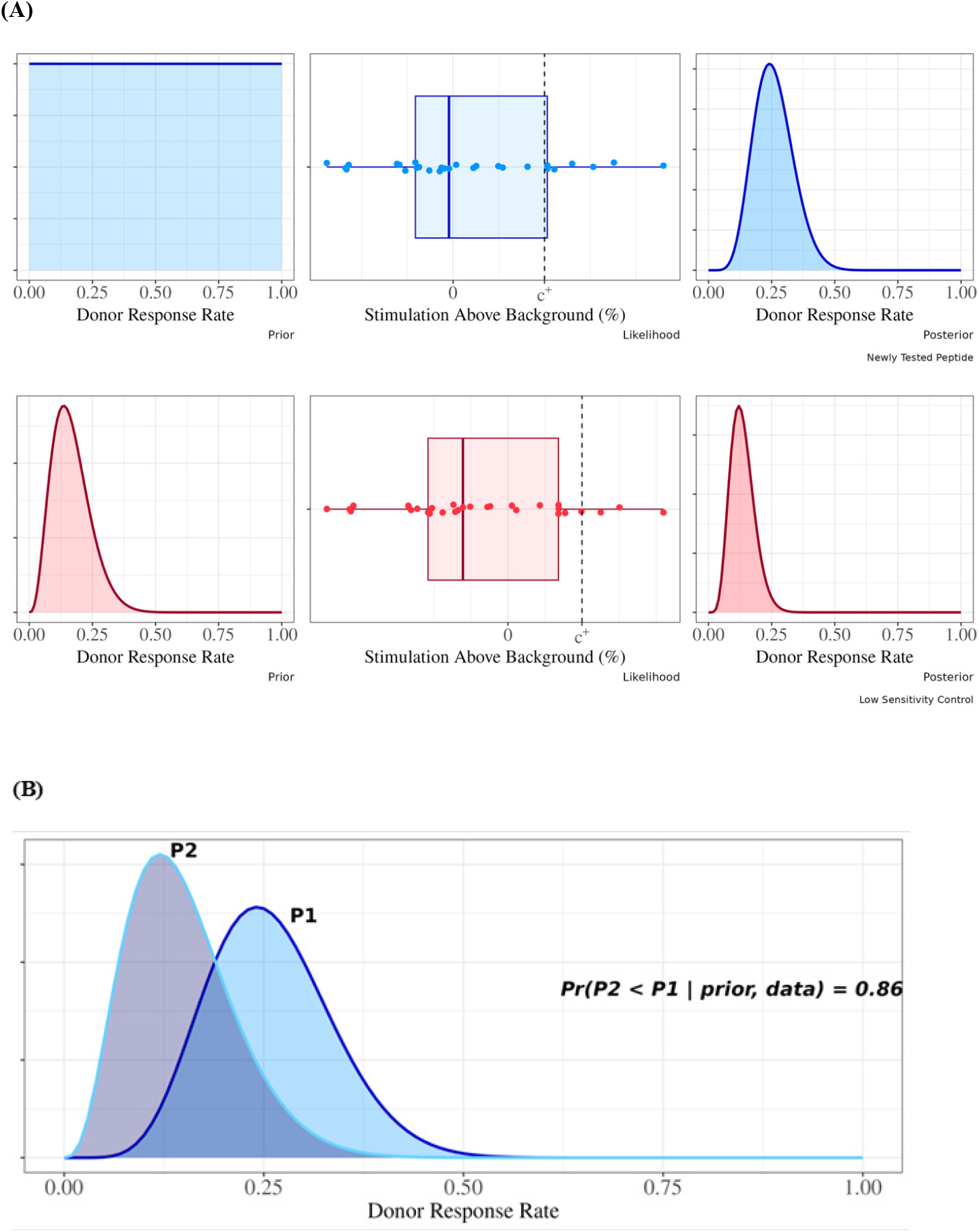
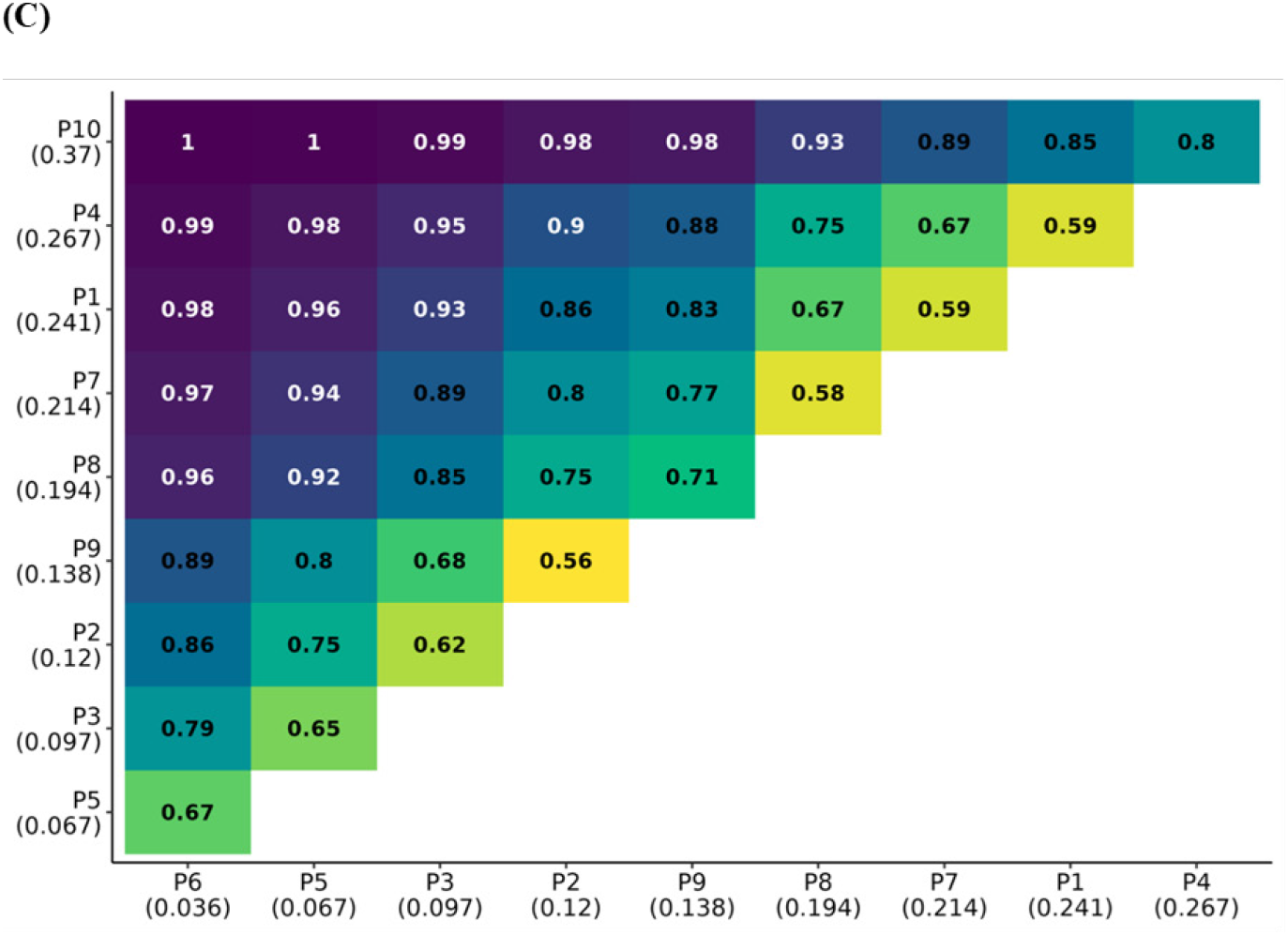
Bayesian analysis of the peptide:donor response rates. (A) Bayesian analysis path incorporating prior donor response rate knowledge, data, and updated posterior belief, for both a newly tested peptide with weak prior knowledge (in blue) and the low sensitivity control with strong prior knowledge (in red). (B) Comparison of Peptide 1 (P1) and Peptide 2 (P2) donor response rate posterior distributions, with probability that P2 has a lower donor response rate than P1. (C) Upper triangular matrix of all (P_i_, P_j_) pairwise comparisons ordered by predicted peptide donor response rate.

## DISCUSSION

The work reported here concerned the characterization of an *in vitro* PBMC:peptide T cell proliferation assay developed to assess the T cell risk of potential CD4+ T cell epitopes present in biologic drug sequences. T cell assays are used across the pharmaceutical industry to contribute to immunogenicity risk assessments and to help select clinical candidates with optimized immunogenicity risk profiles (16).The goal of the present study was to increase confidence in the T cell epitope risk assessment provided by the assay through the establishment of statistically based, data-driven critical assay parameters.

The first step of the assay qualification was to determine donor inclusion criteria for the donor cohort. A key criterium of a donor set is its representativity of the HLA diversity of a population of interest, for instance that of clinical trial or targeted disease populations. Commonly, the donor set representativity is based on the distribution frequency of HLA DR alleles. For instance, HLA diversity can be achieved by selecting a donor set that expresses a combination of HLA II supertypes (a set of 8-9 alleles encompassing the peptide binding specificities of the most prevalent HLA DR alleles) (19, 21-23). Because HLA-DP and HLA-DQ T cell epitopes have been reported for biotherapeutics (24, 25), the donor set representativity would ideally include these alleles. However, such an approach would require access to a larger pool of donors to select from and poses practical issues. Therefore, and similarly to McGill et al. (26) an algorithm was developed to select HLA-typed donors from the PBMC bank that would match the population of interest for the most prevalent HLA DRB alleles.

The second step was to determine the necessary controls to accept or reject assay runs, donors, or data points. Sensitivity controls should match the format of the test articles where feasible, thus single peptides in the PBMC:peptide assay. Given the HLA diversity within donor cohorts, sensitivity control peptides should exhibit broad HLA class II binding to optimize their likelihood of presentation across all donors. Also, sensitivity controls should assess naïve responses if the assay is intended for use during drug design when donors will not have been exposed to the drug. CLIP was chosen as LSC for its binding promiscuity and its endogenous nature, which was expected to translate into low donor response rates (27). As expected, CLIP exhibited low donor response rates in the assay, generally ranging from 0-12% across the studies included in the qualification data set (data not shown). Identification of a HSC, i.e. a single peptide with HLA II promiscuous binding and which would generate naïve positive responses in most donors in a donor set proved difficult. As an alternative, an HLA II-restricted pool of peptides (broad HLA coverage) composed of known immunogenic CD4 T cell epitopes from several common viruses (responses expected in most donors in a donor set) was selected as an HSC (CEFT, see Material and Methods). Nevertheless, experiments efforts to identify a single short peptide HSC are still on-going. On average, HSC generally exhibited a 1-25% donor response rate across the data set (data not shown). Runs with donor response rates for LSC and HSC outside the set range will be rejected. The responses to both controls also serve to monitor assay performance/stability overtime for possible drifts. Because CEFT is composed of multiple epitopes from common viruses, it does not demonstrate that the assay can detect naïve T cell responses to a single peptide but rather validates that no technical issues preventing the development of specific T cell responses occurred. An indirect demonstration of the capability of the assay to detect naïve T cell responses is the observation of positive responses to test peptides during drug development. These and peptide from other sources are under evaluation as potential HSCs. Of note, if applied to assess T cell responses in treated individuals, the demonstration that the assay detects recall responses (current HSC) might suffice.

The assay in the present study was independently developed during a period where several other assays were in development (16). In most cases, these assays were implemented by industry or academic laboratories to identify positive signals that would either indicate why a biologic therapy had a high clinical incidence of ADA, or to identify the relative risk of sequences or candidates prior to clinical trials. Details of analytical qualification were lacking in most cases. An exception is the qualification of DC:CD4+ T cell assay conducted by Siegel et al., where a more detailed analysis of the sources of variability is provided (28). Contributions to assay variance were calculated for treatments, donors, treatment-donor pairs and same donor at different times. The combination of treatment and donors overall accounted for 66% of the assay variability, with small contributions from analytical factors. Although the study was unable to explain 24% of the assay variability, the finding that most variability related to the test molecule and donors demonstrated the suitability of the assay for application to biologic drug immunogenicity assessment. Results from our study are comparable. The combination of peptides, donors, and the peptide donor pairs, labeled as “Non-Method Sources” in Figure 2(A), account for 85.1% of the variability, while the remaining 14.9% is due to “Method Sources” such as run (operator, plate, etc.) and residual (well). The method precision is further increased through the *n=6* well replicates previously mentioned. There are two notable differences between the analyses: First, the study design in Figure 2(A) informs our linear model and fully saturates the degrees of freedom, implying no unexplained variability. The analysis accounts for all variance sources in the data. Second, although both analyses demonstrated the expected impact of diverse donors and treatments / peptides as major variance contributors, our analysis showed donor-to-donor being the dominant source, whereas the treatment differences were the dominant source in Siegel et al. study. In researching this discrepancy, we observed that the CFC (KLH) was included in the DC:CD4+ variance analysis. We deliberately removed the CFC before the analysis as (a) it is not representative of a typically tested peptide for which we want to understand analytical performance and (b) the stimulation data are so high that including those has the potential to skew results and interpretation, even if those data are log-transformed.

It is still common practice for cell-based immunology assays to use a 2-fold signal increase above background as a response positivity threshold (29, 30). However, applying an arbitrary SAB threshold carries the risk of including false negative or false positive donors from the peptide response rate calculation by under or overestimating T cell responses and not integrating the intrinsic assay performance. Previous analysis of low frequency T cell responses highlighted empirical stimulation index thresholds set between 1.5 and 3.0 and recommended to include a statistical analysis to determine the statistically significant differences between non-stimulated and stimulated conditions (31). This is pertinent to the application of naive T cell assays to biologic therapy immunogenicity due to the reported low frequency of T cells with potential to recognize these molecules (32). Our previous work with a dendritic cell assay highlighted the benefits of identifying response with a statistically determined cut-point as a stimulation index of 1.4 (33). With the proliferation of published T cell assays (16), it is currently more realistic to set standards for publishing assay details than standardizing assays themselves. Harmonization of the assays through more detailed descriptions may lead to future standards being set (12).

The number of specific naïve T cell precursors has been established as a sensitive parameter of *in vitro* T cell assays (32) and the number of CD4+ T cells present in the periphery at the time of blood withdrawal varies between donors (34). In the assay, because a fixed number of CD8+ T cell-depleted PBMCs (as opposed to CD4+ T cells) is distributed, it would be possible that for some donors with low CD4+ T cell counts, the number of CD4+ T cells surveyed in the assay is insufficient and leads to false negative responses. We found no statistically significant difference in donor response frequencies to HSC and LSC when the number of total cells distributed was corrected for the CD4+ T cell content (data not shown). Therefore, the protocol does not include the correction, which simplifies the assay workflow.

Human primary cell-based *in vitro* assays are resource intensive, low to medium throughput, lengthy for most of them, making them sometimes unsuitable for fast-paced early drug development. To address this drawback, we maximized the use of automation from donor sample retrieving to data delivery. This includes donor samples retrieving from liquid nitrogen storage by a fully automated freezer on reception of the donor set list generated by the HLA donor selection algorithm; full automation of the flow cytometry analysis immunostaining step by means of the design and built of a fully automated system (35); development of an analysis webtool that integrates all data quality checks and generate the final donor response rate graph within 2 seconds. We achieved a turnaround of 4 weeks for 40 donors, 50 peptides. A complementary approach is the development of in silico tools to progressively replace experimental work time. Such tools encompass algorithms to predict peptide affinity for HLA II molecules, peptide presentation by antigen presenting cells and T cell responses to peptides have been developed as an alternative to *in vitro* peptide HLA binding experiments, MHC-associated peptide proteomics (MAPPs) and *in vitro* T cell assays, respectively (36-38). Mathematical models that can predict protein therapeutic immunogenicity and its possible impact on drug exposure have also been developed and providing adequate validation, might be applied to clinical candidate selection (39-41). Since several input data originate from *in vitro* assay, their use faces the same time-constraints limitations.

## CONCLUSION AND OUTLOOK

By applying a rigorous statistical path and interrogating each parameter of the assay that could contribute to false positive, false negative results, erroneous calculation of the end results and data misinterpretation, the confidence in the output of the PBMC assay was maximized. The work presented here concerns a specific T cell assay that is designed to contribute to biologic drug design by reducing potential T cell epitope content of selected clinical candidates. Whilst different formats and different T cell assay protocols are used across labs, comparisons and interpretation of published data are difficult to perform with a high degree of confidence (12). A way to harmonize understanding of the value of T cell assays for immunogenicity assessment is to standardize reporting practices. Further harmonization include on-going efforts to generate reference sensitivity control standards (for instance several companies collaborate with the Health and Environmental Sciences Institute Immuno-Safety Technical Committee and the National Institute for Biological Standards and Control, UK, to produce a reference panel of biotherapeutics and with the FDA and the National Institute of Standards and Technology, US, to build a reference panel for assays that assess innate immune response modulating impurities (IIRMIs). Meanwhile, it is essential to qualify NCIRA assays to enhance confidence in data quality and minimize the risk of misinterpretation. This step will promote more consistent integration of NICRA assay results in support of immunogenicity mitigation strategies for regulatory submissions.

## ABBREVIATIONS

BRC: Baseline response control
CFC: Cell functionality control
HSC: High sensitivity control
LSC: Low sensitivity control
MAPPs: MHC-associated peptide proteomics
SAB: Stimulation above background

## ACKNOWLEDGMENT

The authors thank Robert Webster and Eve Pickering for critical review of the manuscript.

## CONFLICT OF INTEREST

All authors were employed by Pfizer at the time of the study. JC and TPH are current employees of EpiVax and Quasor respectively.

## REFERENCES

1. Karle AC, Wrobel MB, Koepke S, Gutknecht M, Gottlieb S, Christen B, et al. Anti-brolucizumab immune response as one prerequisite for rare retinal vasculitis/retinal vascular occlusion adverse events. Sci Transl Med. 2023;15(681):eabq5241.

2. Banugaria SG, Prater SN, Ng YK, Kobori JA, Finkel RS, Ladda RL, et al. The impact of antibodies on clinical outcomes in diseases treated with therapeutic protein: lessons learned from infantile Pompe disease. Genet Med. 2011;13(8):729–36. doi: 10.1097/GIM.0b013e3182174703.

3. Prado MS, Bendtzen K, Andrade LEC. Biological anti-TNF drugs: immunogenicity underlying treatment failure and adverse events. Expert Opin Drug Metab Toxicol. 2017;13(9):985–95.

4. Wills CA, Drago D, Pietrusko RG. Clinical holds for cell and gene therapy trials: Risks, impact, and lessons learned. Mol Ther Methods Clin Dev. 2023;31:101125.

5. Immunogenicity Assessment for Therapeutic Protein Products. Guidance for Industry. https://www.fda.gov/regulatory-information/search-fda-guidance-documents/immunogenicity-assessment-therapeutic-protein-products. zU.S. Food and Drug Administration. 2014.

6. Guideline on immunogenicity assessment of therapeutic proteins. European Medicines Agency. https://www.ema.europa.eu/en/documents/scientific-guideline/guideline-immunogenicity-assessment-therapeutic-proteins-revision-1_en.pdf. 2017.

7. Tourdot S, Hickling TP. Nonclinical immunogenicity risk assessment of therapeutic proteins. Bioanalysis. 2019;11(17):1631–43.

8. Rosenberg AS, Sauna ZE. Immunogenicity assessment during the development of protein therapeutics. J Pharm Pharmacol. 2018;70(5):584–94.

9. Jawa V, Terry F, Gokemeijer J, Mitra-Kaushik S, Roberts BJ, Tourdot S, et al. T-Cell Dependent Immunogenicity of Protein Therapeutics Pre-clinical Assessment and Mitigation–Updated Consensus and Review 2020. Frontiers in Immunology. 2020;11(1301).

10. Yang TY, Braun M, Lembke W, McBlane F, Kamerud J, DeWall S, et al. Immunogenicity assessment of AAV-based gene therapies: An IQ consortium industry white paper. Mol Ther Methods Clin Dev. 2022;26:471–94.

11. Gokemeijer J, Balasubramanian N, Ogasawara K, Grudzinska-Goebel J, Upreti VV, Mody H, et al. An IQ Consortium Perspective on Best Practices for Bioanalytical and Immunogenicity Assessment Aspects of CAR-T and TCR-T Cellular Therapies Development. Clin Pharmacol Ther. 2024;115(2):188–200.

12. Ducret A, Ackaert C, Bessa J, Bunce C, Hickling T, Jawa V, et al. Assay format diversity in pre-clinical immunogenicity risk assessment: Toward a possible harmonization of antigenicity assays. MAbs. 2022;14(1):1993522.

13. Gokemeijer J, Wen Y, Jawa V, Mitra-Kaushik S, Chung S, Goggins A, et al. Survey Outcome on Immunogenicity Risk Assessment Tools for Biotherapeutics: an Insight into Consensus on Methods, Application, and Utility in Drug Development. AAPS J. 2023;25(4):55.

14. Siegel M, Steiner G, Franssen LC, Carratu F, Herron J, Hartman K, et al. Validation of a Dendritic Cell and CD4+ T Cell Restimulation Assay Contributing to the Immunogenicity Risk Evaluation of Biotherapeutics. Pharmaceutics. 2022;14(12).

15. Wickramarachchi D, Steeno G, You Z, Shaik S, Lepsy C, Xue L. Fit-for-Purpose Validation and Establishment of Assay Acceptance and Reporting Criteria of Dendritic Cell Activation Assay Contributing to the Assessment of Immunogenicity Risk. The AAPS Journal. 2020;22(5):114.

16. Tourdot S, Karle AC, Rosenbaum M, Ackaert C, Le Vu P, Gutknecht M, et al. T cell assays for non-clinical immunogenicity risk assessment: best practices recommended by the European Immunogenicity Platform. Front Immunol. 2025;16(In print):1723110.

17. Riberdy JM, Newcomb JR, Surman MJ, Barbosat JA, Cresswell P. HLA-DR molecules from an antigen-processing mutant cell line are associated with invariant chain peptides. Nature. 1992;360(6403):474–7.

18. McCulloch CE, Searle SR. Generalized, Linear, and Mixed Models. New York: Wiley and Sons; 2001.

19. Greenbaum J, Sidney J, Chung J, Brander C, Peters B, Sette A. Functional classification of class II human leukocyte antigen (HLA) molecules reveals seven different supertypes and a surprising degree of repertoire sharing across supertypes. Immunogenetics. 2011;63(6):325–35.

20. Kruschke JK. (2015). Doing Bayesian data analysis: a tutorial with R, JAGS, and Stan. Amsterdam: Academic Press

21. Paul S, Lindestam Arlehamn CS, Scriba TJ, Dillon MB, Oseroff C, Hinz D, et al. Development and validation of a broad scheme for prediction of HLA class II restricted T cell epitopes. J Immunol Methods. 2015;422:28–34.

22. Lund O, Nielsen M, Kesmir C, Petersen AG, Lundegaard C, Worning P, et al. Definition of supertypes for HLA molecules using clustering of specificity matrices. Immunogenetics. 2004;55(12):797–810.

23. Castelli FA, Buhot C, Sanson A, Zarour H, Pouvelle-Moratille S, Nonn C, et al. HLA-DP4, the most frequent HLA II molecule, defines a new supertype of peptide-binding specificity. J Immunol. 2002;169(12):6928–34.

24. Peyron I, Hartholt RB, Pedró-Cos L, van Alphen F, Brinke AT, Lardy N, et al. Comparative profiling of HLA-DR and HLA-DQ associated factor VIII peptides presented by monocyte-derived dendritic cells. Haematologica. 2018;103(1):172–8.

25. Hartman K, Steiner G, Siegel M, Looney CM, Hickling TP, Bray-French K, et al. Expanding the MAPPs Assay to Accommodate MHC-II Pan Receptors for Improved Predictability of Potential T Cell Epitopes. Biology (Basel). 2023;12(9).

26. McGill JR, Yogurtcu ON, Verthelyi D, Yang H, Sauna Z. SampPick: Selection of a Cohort of Subjects Matching a Population HLA Distribution. Front Immunol. 2019;10:2894.

27. Sette A, Southwood S, Miller J, Appella E. Binding of major histocompatibility complex class II to the invariant chain-derived peptide, CLIP, is regulated by allelic polymorphism in class II. J Exp Med. 1995;181(2):677–83.

28. Siegel M, Padamsey A, Bolender AL, Hargreaves P, Fraidling J, Ducret A, et al. Development and characterization of dendritic cell internalization and activation assays contributing to the immunogenicity risk evaluation of biotherapeutics. Front Immunol. 2024;15:1406804.

29. Lee Y, Tarke A, Grifoni A. In-depth characterization of T cell responses with a combined Activation-Induced Marker (AIM) and Intracellular Cytokine Staining (ICS) assay. Oxf Open Immunol. 2024;5(1):iqae014.

30. Ogese MO, Watkinson J, Lister A, Faulkner L, Gibson A, Hillegas A, et al. Development of an Improved T-cell Assay to Assess the Intrinsic Immunogenicity of Haptenic Compounds. Toxicol Sci. 2020;175(2):266–78.

31. Di Blasi D, Claessen I, Turksma AW, van Beek J, Ten Brinke A. Guidelines for analysis of low-frequency antigen-specific T cell results: Dye-based proliferation assay vs (3)H-thymidine incorporation. J Immunol Methods. 2020;487:112907.

32. Delluc S, Ravot G, Maillere B. Quantitative analysis of the CD4 T-cell repertoire specific to therapeutic antibodies in healthy donors. FASEB J. 2011;25(6):2040–8.

33. Wickramarachchi D, Steeno G, You Z, Shaik S, Lepsy C, Xue L. Fit-for-Purpose Validation and Establishment of Assay Acceptance and Reporting Criteria of Dendritic Cell Activation Assay Contributing to the Assessment of Immunogenicity Risk. AAPS J. 2020;22(5):114.

34. Burel JG, Qian Y, Lindestam Arlehamn C, Weiskopf D, Zapardiel-Gonzalo J, Taplitz R, et al. An Integrated Workflow To Assess Technical and Biological Variability of Cell Population Frequencies in Human Peripheral Blood by Flow Cytometry. J Immunol. 2017;198(4):1748–58.

35. Dhule S, Corriveau E, Lepsy C, Tourdot S. Staining Triad: A fully automated and zero-waste flow cytometry staining system fostering the 3R to 4R transition. SLAS Technol. 2025;35:100345.

36. Yan Z, Kim K, Kim H, Ha B, Gambiez A, Bennett J, et al. Next-generation IEDB tools: a platform for epitope prediction and analysis. Nucleic Acids Research. 2024;52(W1):W526–W32.

37. Haghighatlari M, Marze N, Seward R, Ciarla A, Hindin R, Calderini J, et al. HLAIIPred: cross-attention mechanism for modeling the interaction of HLA class II molecules with peptides. Commun Biol. 2025;8(1):1133.

38. Yogurtcu ON, Sauna ZE, McGill JR, Tegenge MA, Yang H. TCPro: an In Silico Risk Assessment Tool for Biotherapeutic Protein Immunogenicity. AAPS J. 2019;21(5):96.

39. Kierzek AM, Hickling TP, Figueroa I, Kalvass JC, Nijsen M, Mohan K, et al. A Quantitative Systems Pharmacology Consortium Approach to Managing Immunogenicity of Therapeutic Proteins. CPT Pharmacometrics Syst Pharmacol. 2019;8(11):773–6.

40. Chen X, Hickling TP, Vicini P. A mechanistic, multiscale mathematical model of immunogenicity for therapeutic proteins: part 2-model applications. CPT Pharmacometrics Syst Pharmacol. 2014;3:e134.

41. Hamuro L, Tirucherai GS, Crawford SM, Nayeem A, Pillutla RC, DeSilva BS, et al. Evaluating a Multiscale Mechanistic Model of the Immune System to Predict Human Immunogenicity for a Biotherapeutic in Phase 1. Aaps j. 2019;21(5):94.

